# BCKDK, a novel hypoxia-responsive kinase that exacerbates cerebral ischemia injury

**DOI:** 10.1101/2025.09.22.677943

**Authors:** Boya Liao, Fang Zhang, Xinyao Yi, Ziyi Zhang, Leigang Jin, Ruby L.C. Hoo, Leiluo Geng, Wei Jia

**Author notes:** Co-correspondence: Ruby L.C. Hoo, Leiluo Geng, and Wei Jia.

## Abstract

Alterations of circulating amino acid profile have been observed in patients with ischemic stroke. However, whether ischemia disrupts amino acid metabolism in the brain tissue and subsequently potentiates cellular stress and cerebral injury have never been explored. Employing a metabolomics approach combined with metabolic flux analysis, impaired catabolism and significant enrichment of branched-chain amino acids (BCAAs) were identified in mouse primary neuron cells upon oxygen-glucose deprivation. Consistently, BCAA catabolism was also damaged in the brain of mouse with acute ischemic stroke, accompanied with suppressed activity of branched-chain alpha-keto acid dehydrogenase (BCKDH) and upregulation of BCKDH kinase (BCKDK). Furthermore, restoration of BCAA catabolism by suppressing BCKDK via pharmacological inhibitor or silencing RNA dramatically alleviated cerebral ischemia injury in mice. Mechanistically, ischemia induces the expression of BCKDK via hypoxia-inducible factor 1α-mediated transcriptional activation and inhibits BCAA conversion into substrates for tricarboxylic acid cycle, contributing to potentiated energy deficiency, glutamate excitotoxicity, and neuronal injury. Collectively, this study identified BCKDK as a novel hypoxia-responsive factor and a promising therapeutic target for cerebral ischemia injury.

## Introduction

Globally, the annual number of strokes increased substantially in the past decades and stroke has been the second-leading cause of death [1]. Ischemic stroke, accounting for more than 80% of stroke incidence, is mainly caused by a sudden blockage of blood flow in the cerebral blood vessels due to a thrombus or embolus [2]. Insufficient supply of oxygen and glucose to the brain parenchyma disrupts oxidative phosphorylation and energy production, resulting in dysfunction of adenosine triphosphate (ATP)-dependent ion pumps and failure of electrochemical gradients in affected neurons [3]. The energy depletion further induces the depolarization of neuron cell membrane and Ca^2+^ overload and triggers the release of excitatory glutamate, which activates multiple glutamate receptors and promotes more uptake of Ca^2+^ in post-synaptic neurons [3]. On one hand, the elevated intracellular Ca^2+^ substantially stimulates aberrant glutamate release and subsequent prolonged activation of glutamate receptors; On the other hand, Ca^2+^ ions induce the production of reactive oxygen species (ROS), endoplasmic reticulum stress, edema, and neuronal inflammation by activating downstream Ca^2+^-dependent enzymes, eventually causing neuron apoptosis, necrosis and death [4]. Ischemia-induced excessive secretion and actions of glutamate, known as glutamate excitotoxicity, plays a cascading effect in the damage of neurons in cerebral ischemia injury [4]. However, the underlying modulatory mechanisms of glutamate excitotoxicity during ischemic stroke are incompletely understood.

Using the metabolomics approaches, a surge of studies reveal the significant change of circulating amino acid profile after ischemic stroke. Many consistent amino acids, including isoleucine, leucine, valine, glycine, lysine, and glutamate, possess potential as diagnostic or prognostic biomarkers for cerebral ischemia injury [5]. Studies have also generated several biomarker panels consisting of serum/plasma amino acids to discriminate ischemic stroke severity and subtypes, facilitating early-stage detection and precision diagnosis of ischemic stroke [6, 7]. However, during the acute ischemic stroke, the alteration of amino acid metabolism in the brain tissue has not been studied. Whether there are some specific amino acid metabolism pathways that can be targeted to treat cerebral ischemia injury warrants to be explored.

Among all the amino acids, branched-chain amino acids (BCAAs), including leucine, isoleucine, and valine, are featured with the aliphatic side-chain with a branch and account for 35% of the essential amino acids in muscle proteins and 40% of the preformed amino acids required by mammals [8]. In the BCAA catabolic pathway, BCAAs are first converted into branched-chain alpha-keto acids (BCKAs) by branched-chain aminotransferase (BCAT) in a reversible reaction, followed with irreversible decarboxylation catalyzed by rate-limiting enzyme branched-chain alpha-keto acid dehydrogenase (BCKDH), and eventually metabolized to acetyl-CoA or succinyl-CoA for oxidation in the tricarboxylic acid (TCA) cycle [9]. The BCKDH is inactivated through phosphorylation of its E1α subunit by BCKDH kinase (BCKDK) and activated through dephosphorylation by the mitochondria-localized protein phosphatase Mg^2+^/Mn^2+^ dependent 1K (PPM1K) [10]. In the present study, we identified cerebral BCAA catabolism was impaired due to upregulation of BCKDK under ischemic stress and BCKDK is a potential therapeutic target for cerebral ischemia injury.

## Result

### Oxygen-glucose deprivation suppresses BCAA catabolism in mouse primary neuron cells

Oxygen-glucose deprivation (OGD) is a well-established method to induce ischemic stress *in vitro* [11]. To investigate the impact of ischemia on the change of intracellular free amino acids (FAAs), mouse primary neuron cells were subjected to OGD for 4 hours, followed with cell harvest and FAA profiling on an ACQUITY UPLC I-Class system coupled with Xevo TQ-XS mass spectrometer (Waters corporation, USA) (Figure 1A). The relative concentrations of 29 amino acids were calculated and compared between OGD and control groups (Figure 1B and Supplementary Table 1). The top five significantly increased amino acids were glutamine (+245.5%, p<0.0001), citrulline (+205.5%, p<0.0001), isoleucine (+159.0%, p<0.0001), methionine (+156.4%, p=0.0001), and leucine (+151.4%, p<0.0001), while the top five significantly decreased amino acids were proline (-78.5%, p<0.0001), aspartic acid (-73.2%, p<0.0001), alanine (-51.9%, p<0.0001), glycine (-47.1%, p=0.0001), and histidine (-39.8%, p<0.0001) under the ischemic stress, suggesting ischemia causes a systemic and complicated metabolic disturbance of amino acids in neuron cells (Figure 1B). Other than isoleucine and leucine, another member belonging to BCAA family, valine, was also significantly elevated during OGD (+63.6%, p=0.0002), leading to an almost 2.1-fold increase of intracellular total BCAA contents (Figure 1C). Since BCAAs cannot be synthesized in the mammalian cells [8], ischemia-induced BCAA accumulation is supposed to be caused by impaired BCAA catabolism in the neuron cells. To quantitatively assess BCAA catabolism flux in primary neuron cells during OGD, ^13^C-labeled leucine was used to determine isotopic enrichment of derived metabolites by gas chromatography-mass spectrometry (Figure 1D). Analysis revealed that OGD increased isotope-labeled leucine and α-ketoisocaproate (KIC) and decreased labeling of the TCA cycle intermediates, including C5-CoA, acetyl-CoA, citrate, and succinate (Figure 1D), suggesting the downstream catabolism of KIC was impaired in the BCAA catabolic pathway by ischemia. Overall, catabolic flux of BCAAs is reduced by ischemia, causing intracellular accumulation of BCAAs in primary neuron cells.

**Figure 1.**
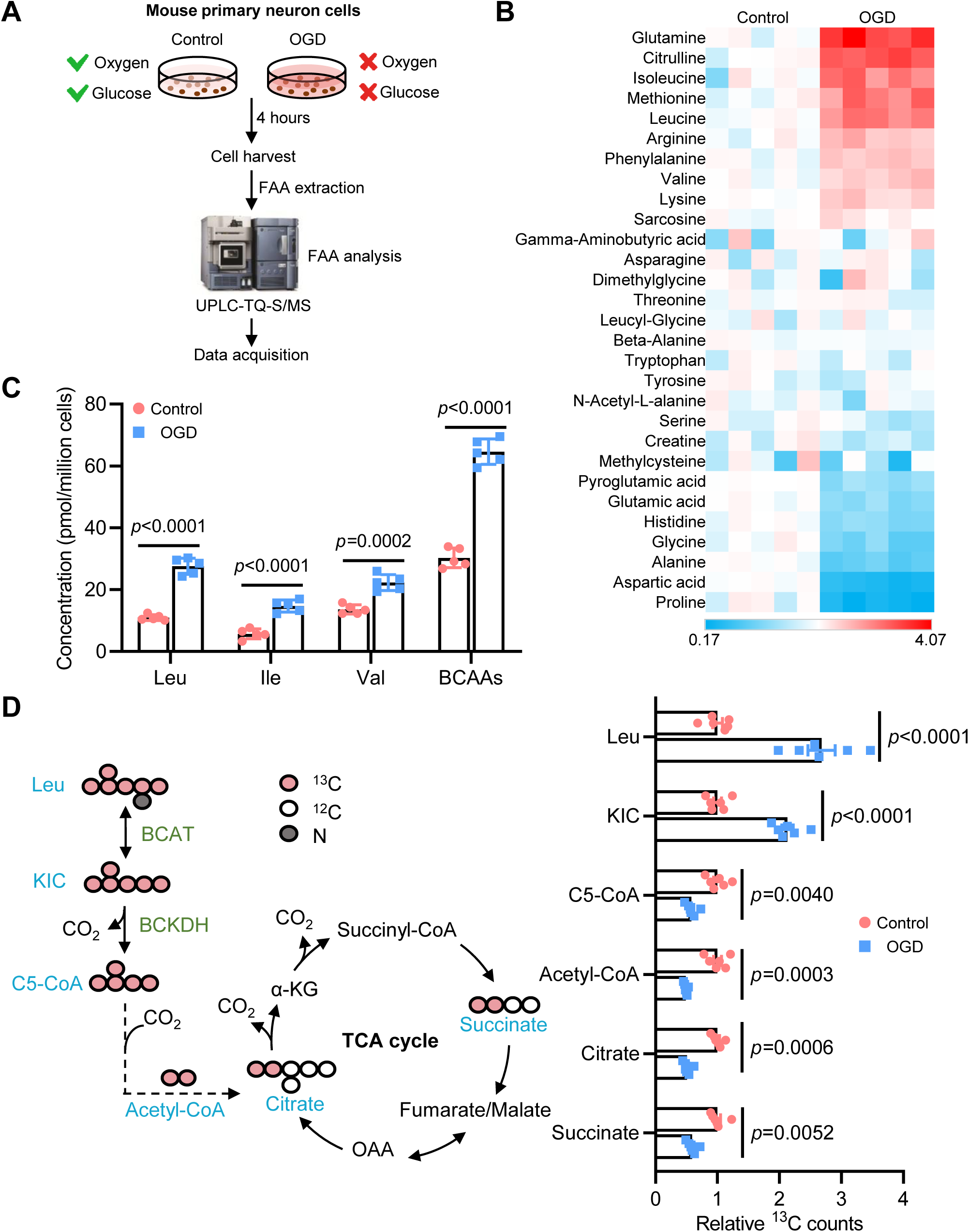
Oxygen-glucose deprivation impairs BCAA catabolism in mouse primary neuron cells. **A.** Experimental design for measurement of intracellular free amino acids (FAAs) in mouse primary neuron cells exposed to oxygen-glucose deprivation (OGD) or normal supply of oxygen and glucose (Control) by using validated ACQUITY UPLC I-Class system coupled with Xevo TQ-S mass spectrometer (UPLC-TQ-S/MS). **B.** Heatmap of the relative levels of 29 FAAs in mouse primary neuron cells under control or OGD condition (n=5). For each amino acid, the average quantity in the control group was set to 1. The full amino acid profile is provided in Supplementary Table 1. **C.** The absolute concentration values of leucine (Leu), isoleucine (Ile), valine (Val), and total BCAAs (Leu+Ile+Val) in mouse primary neuron cells under control or OGD condition (n=5). **D.** Diagram of the ^13^C-labeled leucine catabolic pathway (left panel), in which red circles indicate ^13^C-labeled carbons. ^13^C-labeled metabolites highlighted by blue colour in the pathway were measured in mouse primary neuron cells under control or OGD condition by mass spectrometry (right panel, n=6). KIC, α-ketoisocaproate. Data are presented as mean ± SD. *P* values are determined by multiple t tests with false discovery rate correction (C), or two-way ANOVA followed by Sidak’s multiple comparisons test (D).

### Cerebral ischemia impairs BCAA catabolism in the mouse brain

To investigate the impact of ischemia-reperfusion on cerebral BCAA catabolism, BCAA levels in the brain tissues were monitored in 10-week-old C57BL/6J mice after middle cerebral artery occlusion (MCAO) surgery. Significant elevation of cerebral total BCAA levels was observed in mice with MCAO surgery starting at 2 hours and peaking at 24 hours, followed by gradual decrease to the basal level after 4 days (Figure 2A). Each member of BCAA family, including leucine, isoleucine and valine was found to be simultaneously increased at the early stage and decreased at the late stage of acute ischemic stroke in the brain (Figure 2A). Therefore, consistent with *in vitro* findings (Figure 1), cerebral BCAAs were also accumulated *in vivo* during ischemic stress, suggestive of defective BCAA catabolism induced by ischemia. Considering the activity of BCKDH is a key determinant for BCAA catabolism, which is tightly regulated by the expression of BCKDK and PPM1K (Figure 2B), the expression levels of these critical enzymes related with BCAA catabolism were quantified. BCAT2, responsible for transferring amino groups from BCAA to α-ketoglutarate to produce BCKA and glutamate in the first step of BCAA catabolism, was not changed under ischemic stroke. Inactivation of BCKDH by phosphorylation on Ser293 was significantly enhanced while the total protein level of BCKDH remained stable after cerebral ischemia, accompanied with upregulated expression of BCKDK, which is the kinase catalyzing phosphorylation of the E1α subunit of BCKDH on Ser293. On the contrary, the protein level of PPM1K, phosphatase for activating BCKDH through dephosphorylation, was not altered under ischemia injury (Figure 2B). Consistently, the cerebral mRNA level of *Bckdk* was significantly induced by MCAO surgery, while mRNA levels of other enzymes involved in regulating BCAA catabolism showed no obvious changes (Figure 2C). Furthermore, the enzymatic activity of BCKDH measured by isotope labeling method [12] was remarkably reduced in the brain of mice with MCAO surgery (Figure 2D), in line with the increased phosphorylated isoform of BCKDH (Figure 2B). Collectively, cerebral ischemia-reperfusion injury upregulates the expression of BCKDK and therefore inhibits BCKDH activity and BCAA catabolism, leading to BCAA accumulation in the mouse brain.

**Figure 2.**
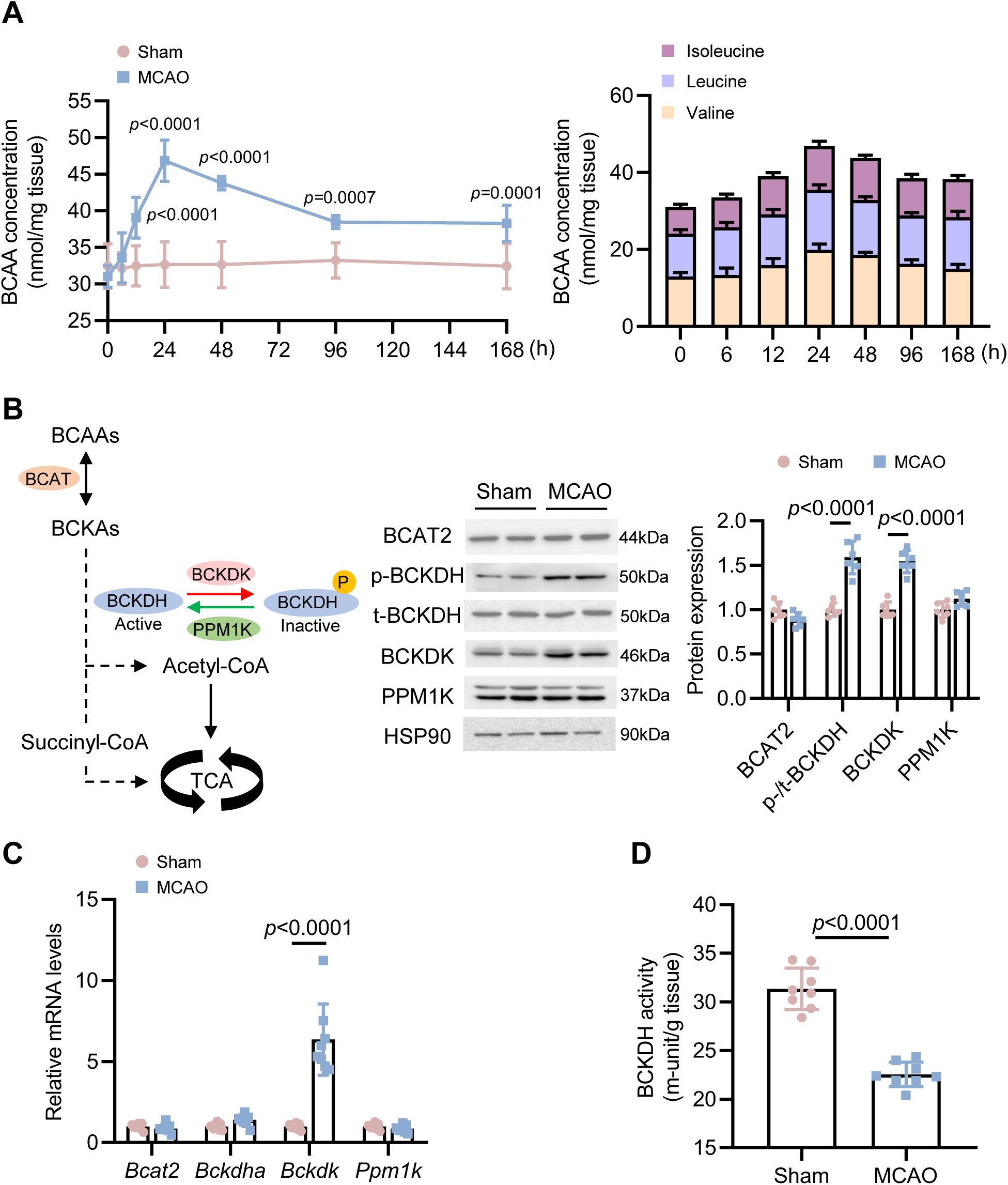
Ischemia-reperfusion injury impairs BCAA catabolism in the mouse brain. 8-week-old male C57BL/6J mice were subjected to MCAO surgery for 1 hour followed with reperfusion for 7 days. The mice were sacrificed for tissue harvest at indicated time points. **A.** The total (left panel) and individual (right panel) BCAA concentrations in infract core of mouse brain at different time points after sham or MCAO surgery (n=8). **B.** BCAA catabolic pathway (left panel), in which the critical enzymes are highlighted with colored background, including BCAT, BCKDK, PPM1K, and BCKDH. Representative western blots (middle panel) and quantitation (right panel) of BCAA catabolic enzymes in infract core of mouse brain after sham or MCAO surgery (n=6). **C.** The mRNA abundance of *Bcat2*, *Bckdha*, *Bckdk*, and *Ppm1k* in the mouse infract core of brain after sham or MCAO surgery normalized to the internal control gene *Gapdh* (n=8). **D.** BCKDH activity was measured using ^13^C-labeled α-ketoisovalerate in infract core of mouse brain after sham or MCAO surgery (n=8). Data are presented as mean ± SD. *P* values are determined by two-tailed unpaired t-test (D) or two-way ANOVA followed by Sidak’s multiple comparisons test (A-C).

### BCKDK is a hypoxia-responsive factor regulated by HIF1α

To verify the regulatory role of ischemia in the expression of enzymes involved in the BCAA catabolism pathway as identified in the brain of mice with MCAO surgery, Neuro2a cells were cultured under the OGD condition for various time periods, followed by examination of the expression of those proteins by immunoblotting. Consistent with *in vivo* findings (Figure 2B), the phosphorylation of BCKDH on Ser293 was gradually enhanced with the increase of OGD period (Figure 3A), accompanied with the upregulation of BCKDK at both protein and mRNA levels (Figure 3A and 3B). Interestingly, the expression of hypoxia-inducible factor 1α (HIF1α), a master transcriptional regulator of cellular and systemic homeostatic response to hypoxia [13], was significantly increased by OGD treatment and closely associated with the expression level of BCKDK (Figure 3A and Supplementary Figure 1), suggesting *Bckdk* is a possible downstream target gene of HIF1α. To test this hypothesis, cobalt chloride (CoCl_2_), a chemical inducer of HIF1α was used to stimulate the HIF1α content in Neuro2a cells, followed by the examination of expression levels of BCKDK. As expected, the protein levels of HIF1α were elevated by CoCl_2_ treatment in a dosage-dependent manner, the same trends of BCKDK expression at both protein and mRNA levels were observed (Figure 3C and 3D). To determine whether hypoxia-induced BCKDK expression was indeed mediated by HIF1α, Neuro2a cells were cultured under OGD conditions with or without treatment of HIF1α selective inhibitor PX-478 for 8 hours. The results showed that inhibition of HIF1α significantly reduced the BCKDK protein and mRNA levels (Figure 3E and 3F). These data suggest that the transcription of *Bckdk* is enhanced by HIF1α during ischemia.

**Figure 3.**
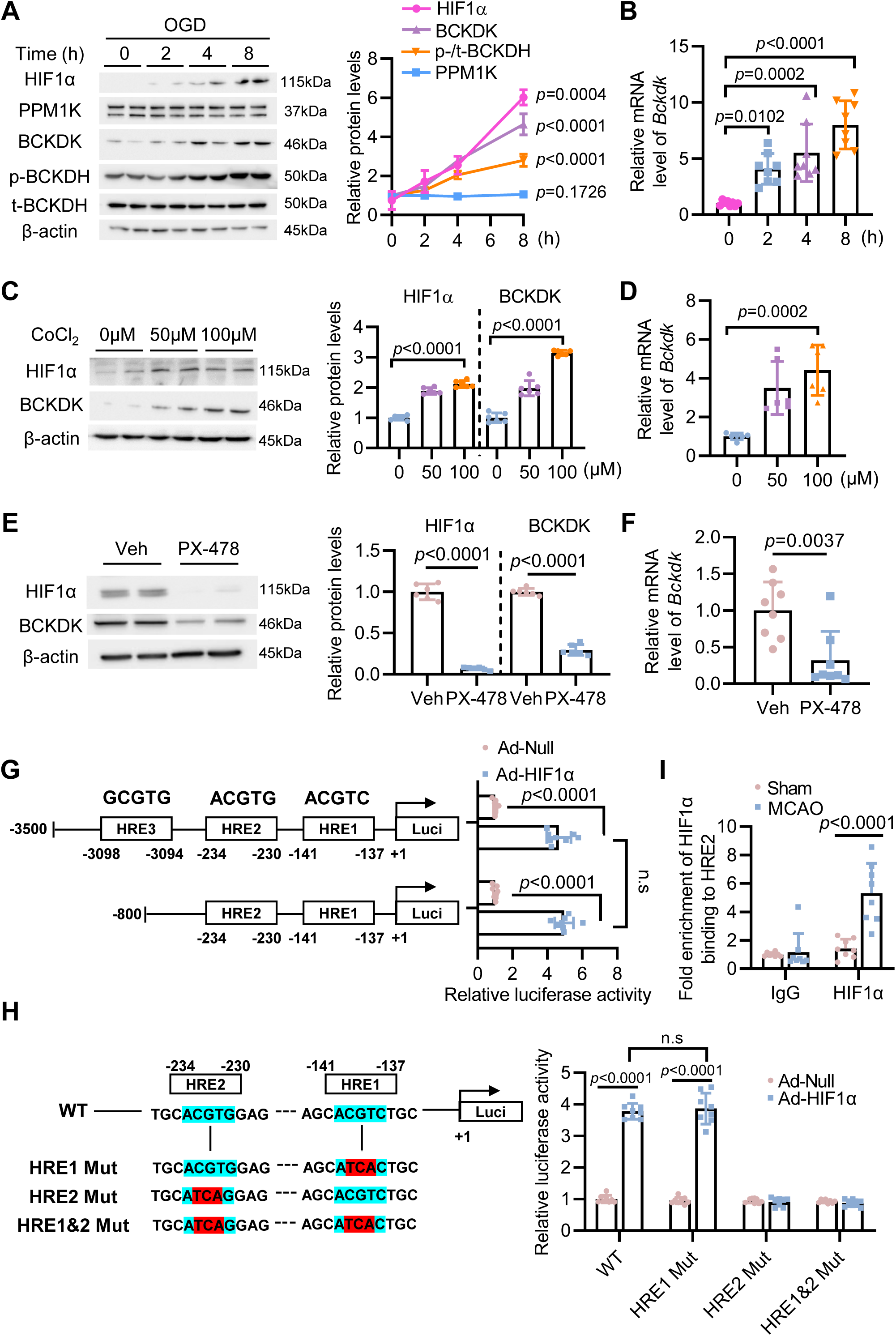
BCKDK is a hypoxia responsive kinase regulated by HIF1α. **A.** Representative immunoblots (left panel) and densitometric quantification of band intensity (right panel, n=4) for HIF1α, BCKDK, PPM1K, p-BCKDH(Ser293)/ t-BCKDH in the mouse primary neuron cells subjected to oxygen-glucose deprivation (OGD) for 0, 2, 4 and 8 hours. β-actin, internal control. B. Relative mRNA levels of *Bckdk* in mouse primary neuron cells under OGD condition for 0, 2, 4, and 8 hours (n=8). C. Representative immunoblots (left panel) and densitometric quantification of band intensity (right panel, n=6) for HIF1α and BCKDK in mouse primary neuron cells treated with CoCl_2_ at indicated concentrations for 4 hours. D. Relative mRNA levels of *Bckdk* in mouse primary neuron cells treated with CoCl_2_ at indicated concentrations for 4 hours (n=6). E. Representative immunoblots (left panel) and densitometric quantification of band intensity (right panel, n=6) for HIF1α and BCKDK in mouse primary neuron cells treated with vehicle (Veh) or HIF1α selective inhibitor PX-478 at 20 µmol/L under OGD condition. β-actin, internal control. F. Relative mRNA levels of *Bckdk* in mouse primary neuron cells treated with Veh or PX-478 under OGD condition (n=8). G. Schematic diagram showing the locations of three predicted hypoxia responsive elements (HRE1-3) within the proximal promoter of murine *Bckdk* gene. Cell-based luciferase reporter assays were performed in mouse Neuro2a cells transfected with luciferase-expressing vectors driven by the full-length *Bckdk* promoter containing HRE1-3 (-3500/+1) or truncated *Bckdk* promoter containing HRE1-2 (-800/+1) followed by infection of adenovirus overexpressing HIF1α (Ad-HIF1α) or Null adenovirus (Ad-Null). The firefly luciferase activity was measured and normalized to Renilla luciferase activity (n=8). n.s, not significant. H. Cell-based luciferase reporter assays were performed in mouse Neuro2a cells transfected with luciferase-expressing vectors driven by the wildtype murine *Bckdk* promoter (WT) or the mutant promoters (HRE1 Mut, HRE2 Mut, HRE1&2 Mut), in which the core motif -CGT-of either HRE1 or HRE2 or both were mutated into -TCA-, together with infection of Ad-HIF1α or Ad-Null. The firefly luciferase activity was measured and normalized to Renilla luciferase activity (n=8). n.s, not significant. I. Binding of HIF1α protein to HRE2 region within the murine *Bckdk* promoter in the brain tissue of mice subjected to sham or MCAO surgery by a ChIP assay (n=8). Data are presented as mean ± SD. *P* values are determined by one-way ANOVA followed by Tukey’s multiple comparisons test (A, B, C, D), two-way ANOVA followed by Tukey’s multiple comparisons test (G, H, I) or two-tailed unpaired t-test (E, F).

To further investigate how HIF1α activates the transcription of *Bckdk* gene, a comprehensive promoter sequence analysis of murine *Bckdk* gene was performed and three potential hypoxia responsive elements (HREs) were identified within the promoter region of the murine *Bckdk* gene, namely HRE1 (-141ACGTC-137), HRE2 (-234ACGTG-230), and HRE3 (-3098GCGTG-3094) (Figure 3G). To determine which HRE site is responsible for *Bckdk* expression, luciferase-reporter constructs containing a murine *Bckdk* promoter fragment spanning from -3500 to -137 (including HRE1 to 3) or from -800 to -137 (HRE1 and HRE2) were transfected into Neuron2a cells, with or without adenovirus-mediated HIF1α overexpression. The reporter activity showed that the lack of HRE3 had little impact on the reporter activity, indicating that HRE3 is not responsible for *Bckdk* transcription by HIF1α (Figure 3G). Next, to determine the contribution of HRE1 and HRE2 to *Bckdk* expression, luciferase-reporter constructs containing a murine *Bckdk* promoter fragment spanning from -800 to -137 (WT), promoters in which either one (HRE1 or HRE2) or both (HRE1&2) core motives of HRE sites were mutated (CGT to TCA) were transfected into Neuron2a cells with or without adenovirus-mediated overexpression of HIF1α (Figure 3H). Mutation of HRE2, but not HRE1, abolished HIF1α-mediated activation of the *Bckdk* promoters, suggesting that HRE2 is essential for the induction of *Bckdk* by HIF1α (Figure 3H). A chromatin immunoprecipitation assay, using specific primers spanning HRE2 site, showed that ischemia significantly increased the binding between HIF1α and the HRE2 site within *Bckdk* promoter in the brain tissue (Figure 3I). Collectively, these data indicate that HIF1α enhances *Bckdk* expression by binding to the HRE2 site within the promoter of *Bckdk*, triggering the transactivation of *Bckdk* during hypoxia.

### Pharmacological inhibition of BCKDK attenuates cerebral ischemia injury in mouse

To investigate whether inhibition of the upregulated BCKDK is able to mitigate cerebral ischemia injury, 8-week-old male C57BL/6J mice were orally administered with BT2 (3,6-dichlorobrenzo[b]thiophene-2-carboxylic acid, 40 mg/Kg/day), a selective small-molecular allosteric inhibitor of BCKDK [14], or vehicle (PBS) for 7 consecutive days after sham or MCAO surgery (Figure 4A). Expectedly, BT2 treatment significantly diminished MCAO-induced phosphorylation of BCKDH protein and reduction of BCKDH activity in the mouse brain (Figure 4B and Supplementary Figure 2A). Consistently, cerebral ischemia-induced BCAA accumulation in the brain tissue and circulation was also alleviated by BT2 administration (Supplementary Figure 2B and 2C). Meanwhile, the brain infarct size was reduced by 32% by BT2 administration in the MCAO groups (Figure 4C). Brain edema was also improved by BT2 treatment as evidenced by significant reduction of cerebral water content in BT2-treated mice after MCAO surgery (Figure 4D). The neurological deficits and survival rate were also improved by BT2 treatment over 7-day monitoring after MCAO surgery (Figure 4E and 4F). Furthermore, BT2 administration also alleviated MCAO-induced upregulation of the activity of Caspase-3, which is the critical enzyme promoting neuronal apoptosis, in the infarct core (Figure 4G). The transcription levels of inflammatory cytokines in the brain tissue were also decreased by BT2 treatment (Figure 4H), suggesting ischemia-induced neuroinflammation was alleviated by BCKCK inhibition. Taken together, these data suggest that pharmacological inhibition of BCKDK promotes BCAA catabolism and alleviates cerebral ischemia injury in mice.

**Figure 4.**
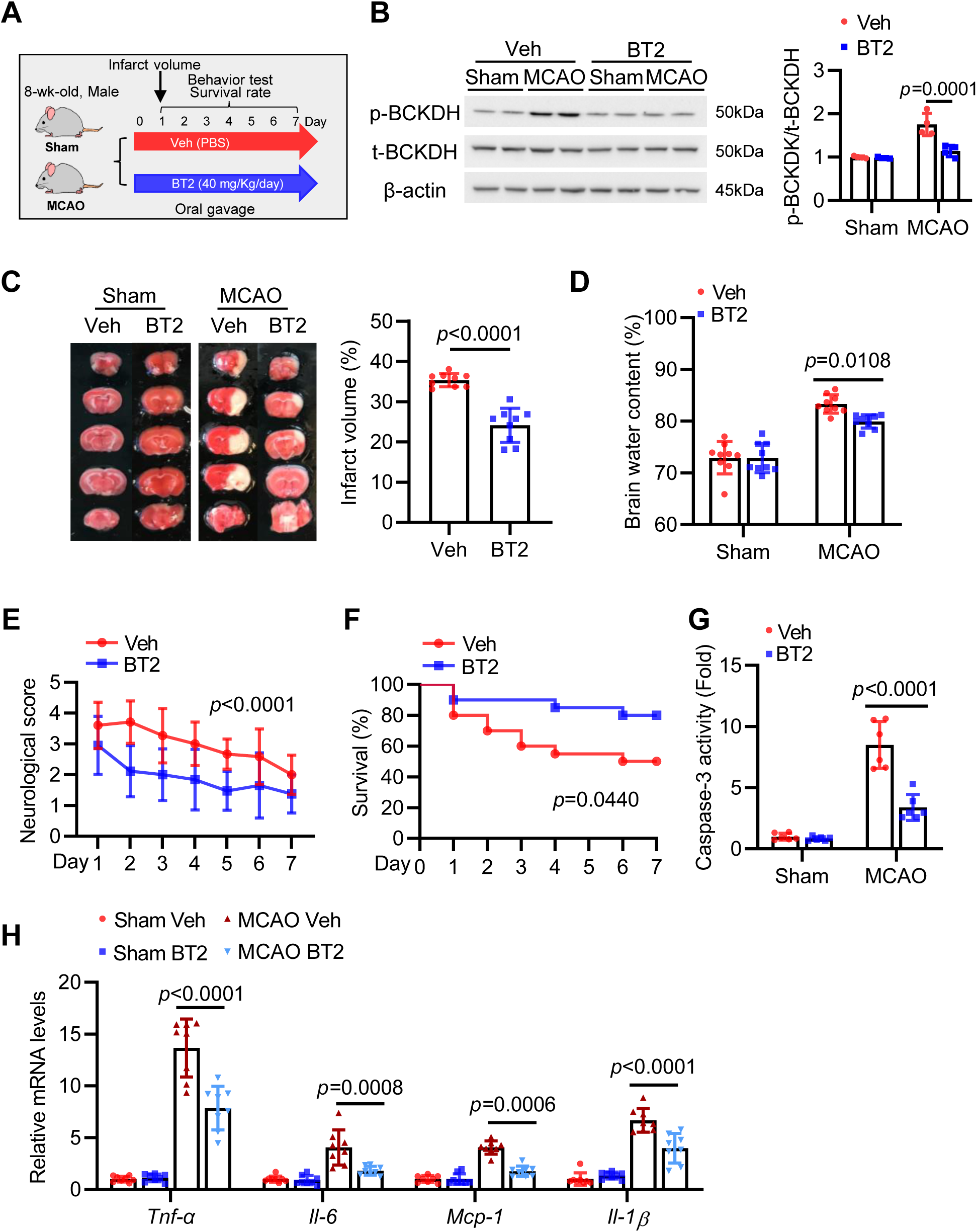
Administration of BT2 mitigates cerebral ischemia injury in mice. **A.** Schematic diagram of animal experimental design. 8-week-old male C57BL/6J mice were subjected to sham or MCAO surgery for 1 hour followed with reperfusion for 7 days. The mice were treated with BT2 (40 mg/Kg body weight per day) or vehicle (PBS) by oral gavage 1 hour post the surgery and every 24 hours for the following six days. **B.** Representative immunoblots and densitometric quantification for p-/t-BCKDH in the infarct core of mouse brain (n=4). β-actin, internal control. **C.** Representative photographs of coronal brain sections stained with TTC 24 hours after sham or MCAO surgery and the quantification of infarct volume in the MCAO groups (n=9). **D.** Percentage of brain water content of mice 24 hours after sham or MCAO surgery (n=9). **E-F.** Neurological score (E) and survival rate (F) of mice during 7-day monitoring after sham or MCAO surgery (n=20 at Day 0). **G.** Caspase-3 activity in the brain tissue of mice 24 hours after sham or MCAO surgery measured by colorimetric assay (n=6). **H.** The mRNA abundance of various inflammatory cytokine genes including *Tumor necrosis factor-α (Tnf-α)*, *Interlukin-6 (Il-6)*, *Monocyte chemoattractant protein-1 (Mcp-1)*, *and Interlukin-10 (Il-10)* in the infarct core of mice treated with vehicle or BT2 24 hours after sham or MCAO surgery (n=8). Data are presented as mean ± SD. *P* values are determined by two-way ANOVA followed by Sidak’s multiple comparisons test (B, D, E, G, H), two-tailed unpaired t-test with Welch’s correction (C), or log-rank test (F).

### BCKDK knockdown by siRNA alleviates cerebral ischemia injury in mouse

To avoid whole-body effects of pharmacological treatment, small interfering RNA (siRNA) was used to obtain cerebral-specific knockdown of BCKDK in mouse through intracerebroventricular (ICV) injection. Several pairs of siRNA targeting BCKDK (siBCKDK) at different coding regions were designed and their efficacy in reducing BCKDK protein expression was assessed in mouse Neuro2a cells by comparing with an siRNA of scrambled sequence as negative control (siNeg). An optimal siBCKDK, which revealed a consistently maximal knockdown of 90% in reducing BCKDK protein (Figure 5A), was selected for the following *in vivo* studies. 8-week-old male C57BL/6J mice were administered with siBCKDK or siNeg by single bilateral ICV injection. 5 days later, these mice were subjected to either sham or MCAO surgery, followed with 7-day monitoring of cerebral ischemia outcomes (Figure 5B). MCAO-induced upregulation of BCKDK and phosphorylation of BCKDH were suppressed by injection of siBCKDK in the mouse brain 24 hours post-stroke (Figure 5C). Meanwhile, cerebral BCKDH activity was improved and the brain BCAA content was reduced by BCKDK knockdown during ischemic stroke (Supplementary Figure 3A and 3B). Unexpectedly, the serum level of BCAAs was also reduced by cerebral BCKDK knockdown (Supplementary Figure 3C), suggesting the brain tissue is one of the important BCAA consumption organs. After 24-hour reperfusion, significantly lower infarct volume and brain edema were observed in mice with cerebral BCKDK knockdown in the MCAO groups (Figure 5D and 5E). Meanwhile, in the following 7-day monitoring, better neurological score and higher survival rate were detected in mice injected with siBCKDK (Figure 5F and 5G). Additionally, the increase in Caspase-3 activity during ischemic stroke was significantly attenuated by knockdown of BCKDK (Figure 5H). Collectively, siRNA-mediated knockdown of BCKDK in the brain tissue showed a similar effect as BT2 treatment in rescuing ischemia-induced cerebral injury in mice.

**Figure 5.**
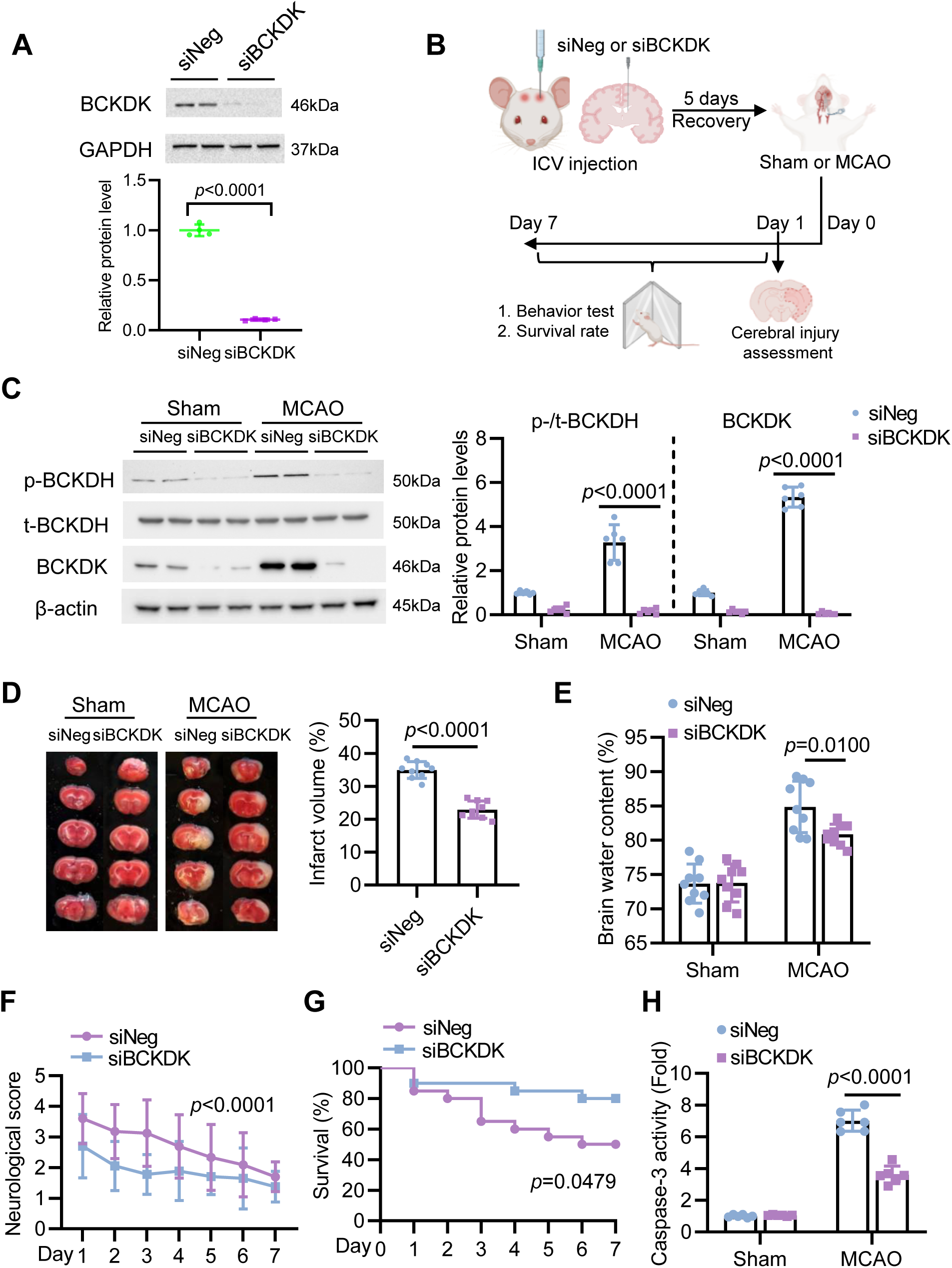
Suppression of cerebral BCKDK by siRNA alleviates ischemic stroke outcome in mouse. **A.** Representative immunoblots and densitometric quantification for BCKDK protein in the mouse Neuro2a cells transfected with siRNA targeting BCKDK (siBCKDK) or siRNA of scrambled sequence as negative control (siNeg) (n=4). GAPDH, internal control. **B.** Schematic representation of animal experimental design. 8-week-old male C57BL/6J mice were administered with siBCKDK or siNeg (235 µg per ventricle) by bilateral intracerebroventricular injection. 5 days later, the mice were subjected to either sham or MCAO surgery, followed by 7-day monitoring of ischemic stroke outcomes, including infarct volume, brain edema, neurological deficits and survival. **C.** Representative immunoblots and densitometric quantification of p-BCKDH, t-BCKDH, and BCKDK proteins in the brain tissues of mice 24 hours after sham or MCAO surgery (n=6). β-actin, internal control. **D.** Representative photographs of coronal brain sections stained with TTC (left) 24 hours after sham or MCAO surgery and the quantification of infarct volume (right) (n=9). **E.** Percentage of brain water content of mice 24 hours after sham or MCAO surgery (n=9). **F-G.** Neurological score (F) and survival rate (G) of mice during 7-day monitoring after sham or MCAO surgery (n=20). **H.** Caspase-3 activity in the brain tissue of mice 24 hours after sham or MCAO surgery (n=6). Data are presented with mean ± SD. *P* values are determined by two-way ANOVA followed by Sidak’s multiple comparisons test (C, E, F, H), two-tailed unpaired t test with Welch’s correction (A, D), or log-rank test (G).

### Impaired BCAA catabolism augments ischemia-induced glutamate excitotoxicity

Next, we sought to investigate the mechanisms whereby how restoration of BCAA catabolism by BCKDK inhibition mitigates cerebral ischemia injury. Ischemic stroke disrupts energy production and causes glutamate excitotoxicity in affected neurons, which plays a catastrophic effect in cerebral ischemia injury [4]. We hypothesized that restoration of BCAA catabolism by BCKDK inhibition may provide more TCA substrates and therefore improve energy production and glutamate excitotoxicity, contributing to alleviation of cerebral ischemia injury. To test this hypothesis, ATP content was measured in freshly isolated brain tissues from mice with/without ischemic stroke under the treatment of BT2 or vehicle as described in Figure 4A. Expectedly, restoration of BCAA catabolism by BT2 treatment significantly increased ATP production and inhibited the generation of ROS in the mouse brain under the MCAO condition (Figure 6A and 6B). Meanwhile, pharmacological inhibition of BCKDK by BT2 significantly decreased the interstitial glutamate levels in the brains of mice after MCAO surgery, suggesting restoration of BCAA catabolism plays a neuroprotective role in cerebral ischemia by reducing glutamate hypersecretion (Figure 6C). Similarly, restoration of cerebral BCAA catabolism by siRNA-mediated BCKDK knockdown also elevated ATP levels, lowered ROS contents, and decreased interstitial glutamate levels in the ischemic brains (Supplementary Figure 4A-4C). When the glutamate concentration increases, it further binds and activates glutamate receptors and causes Ca^2+^ influx, activating intracellular Ca^2+^-dependent enzymes and downstream neuron death pathways [15]. Immunoblot results revealed a significant activation of Ca^2+^-dependent enzymes in the mouse brain by MCAO surgery, as evidenced by enhanced phosphorylation of calcium/calmodulin-dependent protein kinase IV (CaMKIV) and cAMP-response element binding protein (CREB). However, these changes were dramatically inhibited by BT2 treatment (Figure 6D), further supporting the notion that restoration of BCAA catabolism effectively alleviates ischemia-induced glutamate excitotoxicity. To validate the effects of BT2 in rescuing glutamate excitotoxicity caused by ischemia, mouse primary neuron cells were treated with BT2 or vehicle under the normal or OGD condition (Figure 6E). In line with *in vivo* findings, BT2 treatment attenuated OGD-induced impairments of ATP production, excessive secretion of glutamate into the conditional medium, reduction of cell viability, and activation of intracellular Ca^2+^-dependent enzymes (Figure 6F-6I). Thus, these data indicate that restoration of BCAA catabolism protects against cerebral ischemia injury via enhancing ATP production and alleviating glutamate excitotoxicity.

**Figure 6.**
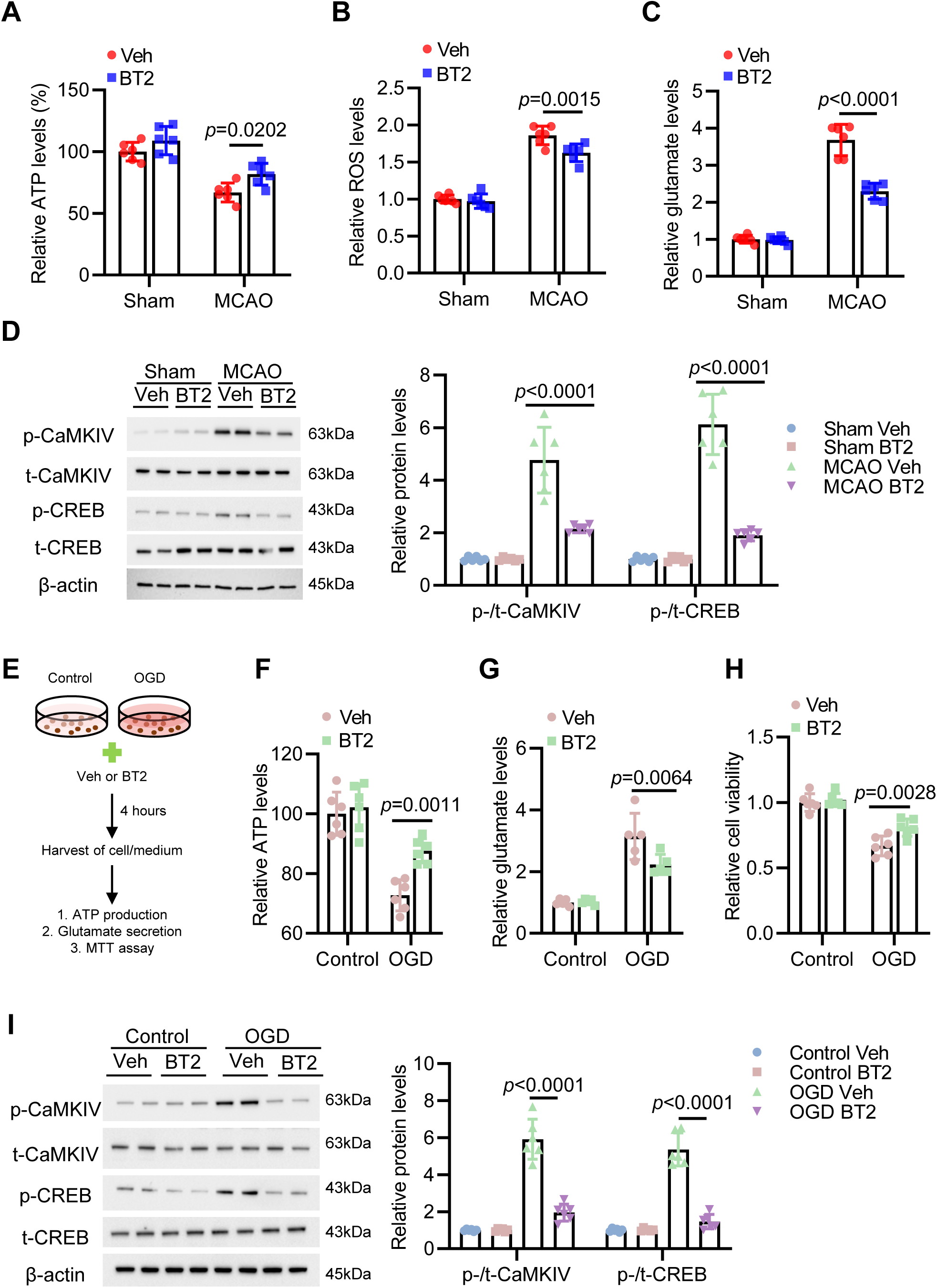
Impaired BCAA catabolism exacerbates glutamate excitotoxicity during ischemia. For panel A-D, the samples were collected from animal experiment as shown in Figure 4A. **A-C.** The relative levels of ATP (**A**), ROS (**B**), and glutamate (**C**) in the brain tissues of mice treated with vehicle or BT2 24 hours after sham or MCAO surgery. (n=6) **D.** Representative immunoblots and densitometric quantification of p-CaMKIV (Thr196/ Thr200), t-CaMKIV, p-CREB (Ser133), and t-CREB in the brain tissues of mice treated with vehicle or BT2 24 hours after sham or MCAO surgery (n=6). β-actin, internal control. **E.** Experimental design for the cell study. Mouse primary neuron cells were subjected to OGD or control condition with the treatment of BT2 or vehicle for 4 hours, followed with measurement of intracellular ATP, glutamate in the medium, and cell viability by MTT assay. **F.** Relative intracellular ATP levels of primary neuron cells cultured under control or OGD condition treated with vehicle or BT2 (n=6). **G.** Relative glutamate concentration in the medium of primary neuron cells cultured under control or OGD condition treated with vehicle or BT2 (n=5). **H.** Relative cell viability in primary neuron cells cultured under control or OGD condition treated with vehicle or BT2 (n=6). **I.** Representative immunoblots and densitometric quantification of p-CaMKIV (Thr196/ Thr200), t-CaMKIV, p-CREB (Ser133), and t-CREB in the primary neuron cells cultured under control or OGD condition treated with vehicle or BT2 (n=6). Data are presented with mean ± SD. *P* values are determined by two-way ANOVA followed by Sidak’s multiple comparisons test (A, B, C, F, G, H) or Tukey’s multiple comparisons test (D, I).

## Discussion

This study demonstrates that cerebral ischemia activates the axis of HIF1α-BCKDK, which inactivates the activity of BCKDH and impairs BCAA catabolism in the brain tissue. Reduced BCAA catabolism contributes to deficiency of TCA substrates and ATP production, leading to enhanced glutamate excitotoxicity and cerebral ischemia injury. Both pharmacological inhibition and siRNA-mediated knockdown of BCKDK are able to restore defective BCAA catabolism and alleviate cerebral ischemia outcomes in mice (Figure 7). Thus, BCKDK was identified as a novel hypoxia-responsive kinase that exacerbates cerebral ischemia injury and represents a promising therapeutic target for ischemic stroke.

**Figure 7.**
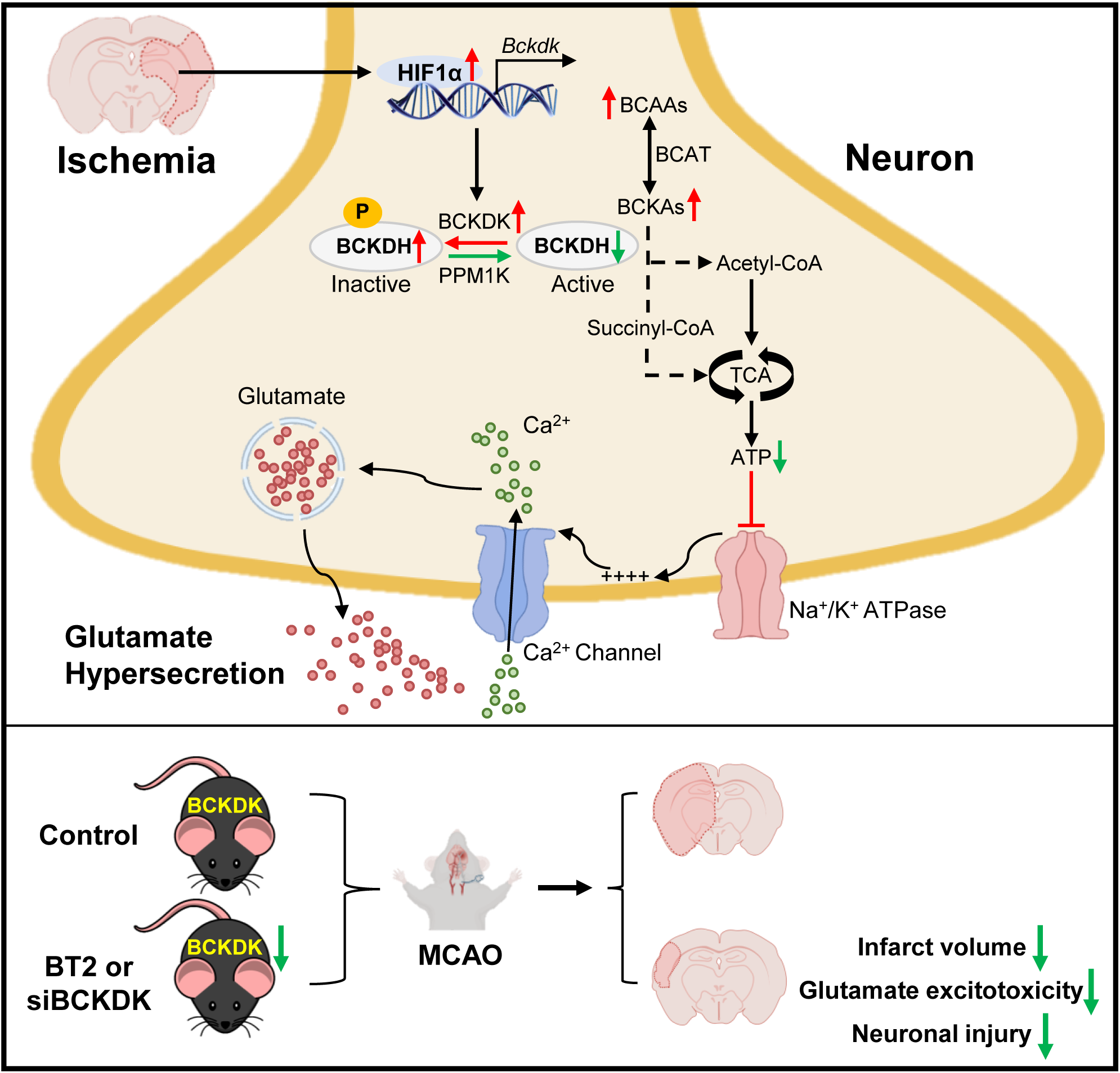
Summary of the findings of this study. Ischemic stroke upregulates the expression of BCKDK via HIF1α in cerebral neurons. Elevated BCKDK phosphorylates and inactivates BCKDH, which is the rate-limiting enzyme in the BCAA catabolic pathway, resulting in impaired BCAA conversion into substrates for TCA cycle and potentiated ATP deficiency, which finally causes glutamate hypersecretion and excitotoxicity as well as enhanced cerebral ischemia injury. Both pharmacological inhibition (BT2 treatment) and siRNA-mediated knockdown of BCKDK mitigate MCAO-induced stroke outcomes in mice, including reduced infarct volume, glutamate excitotoxicity, and neuronal damage, indicating BCKDK is a promising therapeutic target for ischemic stroke. Abbreviations: ATP, adenosine triphosphate; BCAA, branched-chain amino acid; BCAT, branched-chain aminotransferase; BCKA, branched-chain alpha-keto acids; BCKDH, branched-chain alpha-keto acid dehydrogenase; BCKDK, BCKDH kinase; HIF1α, hypoxia inducible factor 1α; MCAO, middle cerebral artery occlusion; Na^+^/K^+^ ATPase, sodium-potassium adenosine triphosphatase; PPM1K, protein phosphatase Mg^2+^/Mn^2+^ dependent 1K; siBCKDK, small interfering RNA targeting BCKDK; TCA, tricarboxylic acid cycle.

A plethora of neurological disorders share a final common deadly pathway known as glutamate excitotoxicity, such as spinal cord injury, stroke, traumatic brain injury, Alzheimer’s disease, Parkinson’s disease, Huntington’s disease, and schizophrenia [16]. If the excessive secretion of excitatory glutamate is failed to be controlled by tight physiological regulation, subsequent prolonged activation of glutamate receptors will form a vicious circle between elevated contents of intracellular Ca^2+^ ions and aberrant glutamate release, aggravating the catastrophic impact on nervous tissues [4]. Our study provides the first clear causal link between BCAA catabolism and glutamate excitotoxicity in cerebral ischemia. First, our *in vitro* study showed that BCAA catabolism was disrupted under OGD conditions, accompanied with reduced ATP production and glutamate hypersecretion in the neuron cells, which were reversed by restoration of BCAA catabolism by BT2 treatment. Second, cerebral glutamate release was enhanced in mice with ischemic stroke induced by MCAO surgery, which was alleviated by restored BCAA catabolism via BT2 treatment or cerebral BCKDK knockdown. Third, the activation of Ca^2+^-dependent enzymes, which is the typical downstream effect of glutamate excitotoxicity [4], was significantly reduced by BT2-stimulated BCAA catabolism in the brain of mouse with ischemic stroke and mouse primary neuron cells under OGD condition. These data revealed that BCAA catabolism modulates ischemia-induced glutamate excitotoxicity in the cerebral neurons and BCAA catabolism pathway may be a potential therapeutic target for multiple neurological diseases related with glutamate excitotoxicity.

Multiple epidemiological studies have confirmed that higher circulating levels of BCAAs are associated with increased risks of obesity and type 2 diabetes (T2D) through comprehensive metabolomic profiling [17, 18]. Considering obesity and T2D are critical predisposing factors for cardiovascular diseases (CVD), recently, the relationship between circulating BCAAs and incident CVD, including stroke, have also been intensively studied, albeit the results are somewhat inconsistent. In a targeted analysis of an unstratified case-cohort study within the Prevención con Dieta Mediterránea (PREDIMED) trial, higher baseline plasma levels of all the three kinds of BCAAs were associated with an elevated risk for stroke [19]. However, in a nested case-control study of the China Kadoorie Biobank, only plasma isoleucine and leucine measured by nuclear magnetic resonance (NMR) spectroscopy were found to be associated with an increased risk of ischemic stroke but not intracerebral hemorrhage [20]. On the contrary, in a prospective cohort of the US Women’s Health Study, neither of the three kinds of plasma BCAAs measured by NMR spectroscopy was found to be significantly associated with incident stroke [21]. The distinct results are possibly caused by the differences in study design, metabolomic profiling approach, study demographics, ethnicity, or stroke type (ischemic or hemorrhagic), compromising the utility of circulating BCAAs as predictive biomarkers for stroke risk. Meanwhile, a two-sample Mendelian randomization study with less complicated confounders and selection bias identified circulating BCAAs were potential causal factors for ischemic stroke, suggesting strategies targeting to reducing systemic BCAAs may be effective in stroke prevention [22]. In the future, a more powerful study, such as randomized control trial, is needed to clarify whether lowering systemic BCAAs is efficacious in reducing the stroke odds via dietary, lifestyle, or medication intervention in high-risk populations.

Despite rigorous studies on the association of baseline circulating BCAAs and incident stroke, the exploration of BCAA metabolism in the brain tissues under acute ischemic stress is scarce. In the present study, we dynamically monitored the BCAAs in the brain tissues of mouse and identified a significant accumulation of BCAAs at 24 hours after the onset of ischemic stroke, which is caused by the impaired BCAA catabolism due to BCKDK upregulation in the mouse brain. Consistently, significant increases of BCAAs in mouse plasma and cerebrospinal fluid were observed at 24 hours post-ischemic stroke compared with respective controls. Interestingly, a previous study performed in rat found the BCAA levels were reduced in plasma and cerebrospinal fluid at 2 hours after the onset of ischemia [23]. These studies suggest circulating BCAA levels may be differentially altered at different time points after the onset of ischemic stroke, the underlying mechanisms of which need to be studied in the future. Remarkably, cerebral knockdown of BCKDK by siRNA was able to reduce the BCAA levels both in the brain and circulation in our mice with ischemic stroke, indicating cerebral BCAA catabolism contributes to the circulating BCAA levels and the brain is also an important organ for BCAA consumption.

Increased BCKDK expression and impaired BCAA catabolism have been observed in different organs under multiple pathophysiological conditions. In *ob/ob* mice and Zucker fatty rats, the protein levels of BCKDK were significantly increased in both liver and adipose tissues compared with lean controls [24]. Similarly, elevated expression of BCKDK was also found in the liver and subcutaneous adipose tissues in humans with severe obesity and positively correlates with steatosis grade, ballooning, and nonalcoholic steatohepatitis [25, 26]. Additionally, BCKDK upregulation was also identified in multiple cancer tissues and predicts tumor aggressiveness and poor prognosis [27–29]. However, the underlying molecular mechanisms whereby how BCKDK is induced in these diseased organs are unclear. In the present study, we found that HIF1α is an important transcriptional factor promoting the expression of BCKDK by binding with the critical HRE region within the proximal promoter of *Bckdk* gene. Since obese adipose and liver tissues as well as multiple tumor tissues are exposed to a hypoxic microenvironment and have elevated HIF1α activity [30–32], HIF1α is likely to play an essential mediating role in modulating BCKDK expression and BCAA catabolism in these pathophysiological conditions. To our knowledge, we are the first to identify *Bckdk* is a target gene of HIF1α and HIF1α-BCKDK axis modulates the BCAA catabolism and hypoxia response.

BT2, a selective, allosteric inhibitor of BCKDK, has been shown to be effective in rescuing insulin resistance and heart diseases in multiple mouse models [33–35], and for the first time, this study has demonstrated its effectiveness in treatment of ischemic stroke. We found BT2 administration via oral gavage enhanced the activity of BCKDH and BCAA catabolism in the brain of mice, contributing to cerebral ATP production and alleviation of glutamate excitotoxicity and cerebral ischemia injury. Thus, BT2 may function as a novel effective therapy for post-ischemic stroke treatment. Previous findings show that BT2 is able to act on multiple organs to promote systemic BCAA catabolism, including heart, liver, kidney, muscle, and adipose tissues [33]. In our study, we found BT2 treatment also enhanced BCAA catabolism in the brain and lowered cerebral BCAA concentration, suggesting BT2 is able to cross the blood-brain barrier and directly act on the cerebral neuron cells. To exclude the interference of other organs, we restored the BCAA catabolism specifically in the ischemic brain by siRNA-mediated knockdown of BCKDK via ICV injection. Significant improvements of cerebral ischemia injury were also obtained by this strategy, indicating BCAA catabolism in local brain tissue indeed modulates the ischemic stroke outcome. Previous studies showed BT2 was well tolerated, with no signs of apparent toxicity, further supporting the therapeutic promise for BCAA-associated cardiometabolic and vascular diseases [36].

In conclusion, we find that BCAA catabolism is a metabolic pathway potently and robustly disrupted in cerebral ischemic neurons, in which HIF1α-induced BCKDK upregulation plays a critical role. Pharmacologic interventions targeting BCKDK inhibition may effectively mitigate ischemic stroke outcomes. Tissue plasminogen activator is the only available pharmacological treatment for acute ischemia stroke up to date but with several limitations, for example, potential bleeding risks and short treatment window [37]. BCKDK inhibition is a potential supplementary therapy to overcome these hurdles.

## Materials and Methods

### Mice

8-week-old male C57BL/6J mice were housed in a temperature-controlled facility (23 °C, 12 h light/dark cycle, 60%-70% humidity) with free access to water and standard chow (#5053, LabDiet). BT2 (#CAS 34576-94-8, Santa Cruz) was administrated by oral gavage at 40 mg/kg body weight at 1 hour after sham or MCAO surgery and once per day during the following 7 consecutive days. Neurological deficit was assessed every day over the 7-day post-surgery monitoring based on Zea-Longa 5-point score criteria: 0 indicates no detectable neurological deficit; 1 indicates failure to extend the forepaw fully; 2 indicates spontaneous contralateral turning; 3 indicates spontaneous contralateral circling; 4 indicates loss of walking ability; and 5 indicates death. The number of living animals in each group was recorded after the sham or MCAO surgery and the survival rate was calculated accordingly. All experimental procedures were approved by the Committee on the Use of Live Animals at The University of Hong Kong and Hong Kong Baptist University.

### Mouse focal ischemia

Mouse focal ischemia was performed by MCAO surgery as previously described [38]. Briefly, mice were anesthetized with 1.5% isoflurane (IsoVet, Chanelle Pharma) in 30%/70% oxygen/nitrous oxide. An MCAO suture made of silicone-wrapped nylon thread (RWD, China) was introduced to the internal carotid artery through external carotid artery and advanced to middle cerebral artery (MCA). Adequate ischemia was confirmed by continuous laser Doppler flowmetry (LDF) with showing at least 30% baseline LDF values during MCAO. After 60min occlusion of MCA, the filament was gently withdrawn to restore the blood flow. The same procedures were preformed except insertion of the filament in the sham groups.

### Evaluation of cerebral infarct volume and water content

The animals underwent transcranial perfusion with saline 1 day after ischemia. Brains were quickly removed, placed within a brain matrix and cut into 8 coronal sections. The sections were then incubated in 2% TTC (2,3,5-Triphenyltetrazolium chloride, #T8877, Sigma) in saline for 10 min at room temperature. Infarction volumes were quantified using the indirect morphometric method with Image J software. To evaluate edema following ischemic stroke, the brain water content was measured 24 hours after MCAO surgery. The left cerebral hemispheres of the mice were collected. Wet tissue samples were weighed immediately after euthanasia. Subsequently, the brain samples were dried in an oven at 110 °C for 48 hours. After drying, the samples were weighed again. The brain water content was determined using the formula: [(wet tissue weight - dry tissue weight) / wet tissue weight] × 100%. All analyses were carried out by operators blinded to conditions.

### Stereotactic injection of siRNA

An *in vitro* validated siRNA targeting BCKDK (siBCKDK, sense: 5’-GUCCGGUACUUC UUGGAUAAATT-3’; antisense: 5’-UUUAUCCAAGAAGUACCGGACTT-3’) and an siRNA of scrambled sequence as negative control (siNeg, sense: 5’-CCUAUGAACGU UAUGACGATT-3’; antisense: 5’-UCGUCAUAACGUUCAUAGGCG-3’) were generated from a commercial biotech company (Genewiz, China). Mice were injected with 470 µg siRNA (235 µg per ventricle) through bilaterally intracerebroventricular injection as previously described [39]. This dose of siRNA was selected for *in vivo* experiments because pilot studies indicated that administration of siRNA up to this dose was well tolerated, and no signs of neurotoxicity (hind-limb paralysis, vocalization, food intake, or neuroanatomical damage) were observed in mice. The mice were anesthetized using Ketamine/Xylazine (70/10 mg/Kg body weight) and kept under anesthesia for 2 hours after injection. Single bilateral intracerebroventricular injections (5 µL per injected side) were performed at 500 nL/min after needle placement at the following coordinates from bregma: -0.2 mm AP, ±0.8 mm mediolateral and -2.5 mm dorsoventral.

### Microdialysis

Microdialysis procedure for cerebral glutamate analysis was performed as previously described [40]. A CX-I probe for brain microdialysis (Eicom, Japan) with specific parameters (4 mm length of the guide cannula, 0.22 mm membrane outer diameter, 0.2 mm membrane inner diameter,1 mm length of membrane, 50 kDa molecular weight cut-off) was inserted into the right striatum of mouse using stereotaxic coordinates (2 mm right lateral from the bregma, 0.5 mm anterior, and 4 mm ventral from the dura). To maintain equilibrium, synthetic cerebrospinal fluid solution containing 125 mM NaCl, 3 mM KCl, 1 mM MgCl_2_, 2 mM CaCl_2_, 26 mM NaHCO_3_, 1.25 mM NaH_2_PO_4_, and 4 mM glucose (pH 7.4) was perfused through the probe at a rate of 1 μL/min. After a 1 hour equilibrium period, microdialysis samples were collected continuously for 4 hours. These samples were then stored at -80 °C for further analysis.

### Cell culture

Mouse primary neuron cultures were generated as previously described [41]. Briefly, the cortical tissues of C57BL/6J mouse embryos at embryonic day 17 were carefully dissected and digested with 0.25% (w/v) Trypsin (#15090046, Gibco) for 15 min. The resulting cells were then plated onto plates coated with 0.1 mg/mL poly-D-lysine solution (#A3890401, Gibco) at a density of 3 × 10^5^ cells/mL in DMEM (#11965084, Life Technology) supplemented with 10% fetal bovine serum (FBS, #10270106, Gibco). After 24 hours, the culture medium was replaced with NeuroBasal medium (#21103049, Invitrogen) supplemented with 2% B-27 (#17504044, Invitrogen), 0.5 mM L-glutamine (#25030149, Gibco), and 1% (v/v) penicillin-streptomycin solution (#15140122, Gibco). The medium was subsequently changed every 3 days and the neuron cells were maintained in a humidified incubator at 37°C and 5% CO_2_ for experiments between 8 to 11 days. Mouse Neuro2a cells were cultured in normal DMEM supplemented with 10% FBS in a humidified incubator at 37°C and 5% CO_2_. To mimic acute ischemia-like condition, cells were cultured with glucose-free DMEM (#11966025, Gibco) pre-equilibrated with 5% CO_2_/ 95% N_2_. The cells were then incubated in a humidified chamber with 5% CO_2_/ 95% N_2_ for indicated periods in each experiment.

### Assay for BCKDH activity

BCKDH activity was measured according to a previous study with some minor modifications [9]. Briefly, 40 mg of brain tissue samples were homogenized using a Qiagen TissueLyser II in 250 μL of pre-cold lysis buffer (50 mM Tris-HCl pH 7.5,150 mM NaCl, 100 μM Na_3_VO_4_, 1% Triton X-100, 1 mM DTT, 3 mM EDTA, 2 mM EGTA, 30 mM NaF, 1 mM α-keto isovalerate, 3% FBS, 1 μM Leupeptin). The lysates were centrifuged at 14,000 g and 4 °C for 15 min and 50 µL supernatant was added into 300 µL of reaction buffer (5.05 M HEPES pH 7.5, 30 mM KPi pH 7.5, 0.4 mM CoA, 3 mM NAD+, 5% FBS, 2 mM Thiamine Pyrophosphate, 2 mM MgCl_2_, 7.8 μM α-keto [^13^C_5_] isovalerate) in a 1.5-mL Eppendorf tube, which was capped and placed at 37 °C for 30 min. The reaction was stopped by adding 100 µL of 70% perchloric acid followed by shaking on an orbital shaker for 1 h. The ^13^CO_2_ was captured using Whatman paper disk containing 20 μL 1 M NaOH at room temperature overnight and counted in 4 mL scintillation fluid (American Biosciences NACS104) with Packard Cobra II Auto Gamma Counter. The BCKDH activity was normalized to the tissue weight.

### Analysis of amino acids

Profiling of intracellular free amino acids in mouse primary neuron cells was conducted using the Q300 Metabolite Assay Kit (Human Metabolomics Institute Inc., China). An ACQUITY ultraperformance liquid chromatography coupled with an XEVO TQ-S mass spectrometry with an ESI source controlled by MassLynx 4.1 software (Waters, USA) was used for the analysis as previously described [42]. Chromatographic separations were performed on an ACQUITY BEH C18 column (1.7 μm, 100 mm × 2.1 mm internal dimensions; Waters, USA). The instrument was operated in positive (POS) and negative (NEG) ion modes. The data were collected with multiple reaction monitor, and the cone and collision energy used the optimized settings from QuanOptimize application manager (Waters, USA). The raw data files were processed using the TMBQ software (v1.0, Human Metabolomics Institute Inc., China) to perform peak integration, calibration, and quantification for each metabolite.

For BCAA analysis in physiological fluid, a reverse phase high-performance liquid chromatography using 1260 Infinity (Agilent Technologies, USA) was adopted as previously described [43]. Amino acid separation and detection were performed by precolumn o-phthalaldehyde derivatization and fluorescent detection (λex = 338 nm, 10-nm bandwidth; λem = 390 nm, 20-nm bandwidth). A gradient elution with mobile phase A (10 mM NaH_2_PO_4_, 10 mM Na_2_B_4_O_7_, 0.5 mM NaN_3_, pH 6.8) and mobile phase B (acetonitrile 45%: methanol 45%: H_2_O 10% V:V:V) was performed with a flow of 1.5 mL/min. The quantification of the amino acid amounts was achieved using calibration curves of external standards of the amino acids of interest with known increasing concentration ranging from 5 to 1,000 µM.

### Analysis of stable isotopic labeling

For isotopic tracer analysis, the cells were washed twice with PBS and changed to fresh medium, in which leucine was replaced with [^13^C_6_]leucine (50 mg/L, #CNLM-281-H, Cambridge Isotope Laboratories). After 90 min, the cells were washed twice with cold PBS to stop metabolic reactions. The cells were lysed and extracted with pre-cold 70% ethanol (v/v) at -80 °C for 30 min and centrifuged at 15,000 g for 20 min at 4 °C to separate soluble and insoluble components. The supernatant was lyophilized and solubilized in water for further gas chromatography-mass spectrometry (GC-MS) analysis according to a previous method [44] using an Agilent 7890 GC equipped with a 30 m DB-35MS capillary column connected to an Agilent 5975B MS operating under electron impact ionization at 70 eV. Data are presented as percentage of labeling of the isotopolog M+X, where M corresponds to the molecular weight of the unlabeled molecule, and X is the number of ^13^C-enriched carbon atoms in the molecule.

### Assays for glutamate, ATP and ROS

Glutamate in conditional medium or microdialysis samples was measured using the Glutamate Assay Kit (#MET-5080, Cell Biolabs). 50 μL of glutamate standard or each sample was added into the wells of a 96 well plate, followed with adding 200 μL of reaction mix and mixing thoroughly. The mixture was incubated at room temperature for 60 min on an orbital shaker and optimal density was detected at 450 nm using a microplate reader.

ATP content in primary neuron cells and fresh brain tissues were measured by an ATP Assay Kit (#ab83355, Abcam). 1×10^6^ cells were harvested, washed with PBS, and then resuspended in 100 μL of ATP assay buffer. Cells were quickly homogenized by pipetting up and down and then centrifuged at 13,000 g at 4 °C for 5 min to remove any insoluble material. As for tissue samples, 10 mg of fresh tissues were washed with cold PBS and homogenized in 100 μL of ATP assay buffer with a Dounce homogenizer, followed with centrifugation at 13,000 g at 4 °C for 15 min. The supernatant was collected and deproteinized by using a Deproteinizing Sample Preparation Kit (#ab204708, Abcam). The supernatants were then collected and incubated with reaction mix at room temperature for 30 min protected from light. Optimal density was detected at 570 nm using a microplate reader.

ROS in freshly-isolated brain tissues was detected by a ROS Fluorometric Assay Kit (#E-BC-K138-F, Elabscience). In brief, the brain tissues were washed with pre-cold PBS and cut into 1 mm^3^ pieces and digested with 0.25% (w/v) Trypsin in 37 °C water bath for 30 min, followed with digestion stopping by adding 10% FBS and filtration through 40 μm cell strainers to prepare single-cell suspension solution. 5 μM DCFH-DA was added into the cell suspension, followed with incubation at 37 °C for 1 hour and centrifugation at 1,000 g for 8 min to collect the cells. The collected cells were resuspended in PBS and the fluorescence was detected on a fluorescence microplate reader with the excitation wavelength of 500 nm and the emission wavelength of 525 nm.

### MTT Assay

The MTT [3-(4,5-dimethylthiazol-2-yl)-2,5-diapehnyltetrazolium bromide] assay (#ab211091, Abcam) was performed to evaluate the cell viability. 5 × 10^4^ cells were seeded in 100 μL culture medium into 96-well tissue culture microplate, followed with incubation for 24 h at 37 °C and 5% CO_2_. The MTT reagent was added into the medium (0.5 mg/mL) and incubated for 3 hours at 37 °C and 5% CO2. After incubation, 100 μL of MTT solvent was added to each well followed with overnight incubation. The absorbance of the formazan product was read by a microplate reader at 570 nm.

### Luciferase assay

The DNA fragments of full-length (-3500/+1), truncated (-800/+1), and HRE-mutated promoter sequences of murine *Bckdk* gene were synthesized by a commercial company (Genewiz, China) and subcloned into pGL3-Basic vector (#E1751, Promega), followed with double confirmation by automatic DNA sequencing. Mouse Neuro2a cells were seeded in 24-well plates and transfected with a firefly reporter vector (0.2 mg) and Renilla reporter vector (0.01 mg) using Lipofectamine™ LTX Reagent with PLUS™ Reagent (#A12621, Invitrogen) when cells had grown up to 90% confluence. After transfection for 24h, cells were infected with adenovirus encoding Myc-tagged HIF1α (Ad-HIF1α) or null adenovirus (Ad-Null) at 100 MOI for 24 h [45]. Finally, the cells were solubilized in a lysis buffer (#E1500, Promega) and luciferase activity was measured with CLARIOstar 0430 Microplate Reader with CLARIOstar Software v.5.01 R2 from BMG LABTECH. The ratio of firefly and Renilla luciferase reading was calculated to indicate the ability of HIF1α to activate HRE-mediated luciferase expression.

### ChIP assay

ChIP assay was performed using a High-sensitivity ChIP Kit (#ab185913, Abcam). Briefly, 40 mg of brain tissue was cut into small pieces of 1 mm^3^ and cross-linked with 1% formaldehyde at room temperature for 20 minutes, which was stopped by adding 1.25 M glycine. The tissue pieces were washed with cold PBS and homogenized with a Dounce homogenizer in working lysis buffer. Cross-linked chromatin was sonicated with a probe microtip (30% power and 8 pulses of 10 second on and 50 second rest on ice between each pulse) and precipitated with non-immune rabbit IgG or rabbit monoclonal IgG against HIF1α (#ab308433, Abcam). Quantitative real-time PCR was performed to measure the amount of bound DNA using primers covering HRE2 region within murine *Bckdk* promoter (Forward primer: CCAACCTCAGCCTGATTGTAT; Reverse primer: CATCTATTGGAGTGGTGGAAAGA).

### Western blot

Protein samples (20-60 μg/well) were loaded onto 8-12% Tris-glycine gels, followed with electrophoresis and protein transfer to PVDF membranes (#88520, Thermo Scientific). The membranes with protein were then blocked in Tris-buffered saline containing 5% non-fat milk for 60 min at room temperature and then incubated overnight at 4 °C with primary antibodies against BCAT2 (#ab95976, Abcam, 1:3,000), BCKDH (#ab126173, Abcam, 1:3,000), p-BCKDH (#ab302504, Abcam, 1:3,000), BCKDK (#ab128935, Abcam, 1:3,000), CaMKIV (#4032, Cell Signaling Technology, 1:1,000), p-CaMKIV (#PA5-37504, ThermoFisher, 1:3000), CREB (#4820, Cell Signaling Technology, 1:1,000), p-CREB (#87G3, Cell Signaling Technology, 1:1,000), HIF1α (#ab308433, Abcam), PPM1K (#ab135286, Abcam, 1:2,000), β-actin (#4967, Cell Signaling Technology, 1:3,000), HSP90 (#4874, Cell Signaling Technology, 1:3,000) or GAPDH (#5174, Cell Signaling Technology, 1:3,000). After intensive washing, the membranes were incubated with corresponding HRP-conjugated secondary antibodies (rabbit, #7074, Cell Signaling Technology, 1:3,000; mouse, #7076, Cell Signaling Technology, 1:3,000; and goat, #ab6741, Abcam, 1:5,000). The protein bands were visualized with enhanced chemiluminescence reagents (Clarity Western ECL Substrate, Bio-Rad 1705061) under the ChemiDocTM MP Imaging System equipped with Image Lab Touch Software 3.0 (Bio-Rad) and quantified using the NIH ImageJ software.

### Quantitative real-time PCR

Total RNA was extracted from cells or tissues using Trizol reagent (#RR037A, Takara). The extracted RNA (400ng as template) was then converted into complementary DNA through reverse transcription, utilizing the PrimeScript RT reagent kit (#RR420D, Takara). For the subsequent analysis of quantitative real-time PCR, SYBR Green One-Step Kit (#1725150, Bio-Rad) was utilized on the Applied Biosystems Prism 7000 Sequence Detection System with StepOnePlus TM Software 2.3 (Applied Biosystems, USA). The expression of target gene was normalized to *18S rRNA*. The primer sequences employed in this experiment can be found in Supplementary Table 2.

### Statistical analysis

Quantitative data were displayed as mean ± SD (standard deviation). Data were calculated and analyzed using GraphPad (Prism v.8.4.3). Statistical significance was determined by two-tailed unpaired Student’s t-test with or without Welch’s correction (for comparison of two experimental conditions) or one/two-way ANOVA followed by Tukey’s or Sidak’s multiple comparisons test (for comparison of three or more experimental conditions). Survival rate was analyzed by the log-rank test. A *p* value less than 0.05 was considered statistically significant.

## Acknowledgement

This study was supported by the National Natural Science Foundation of China (32000816), Guangdong Natural Science Fund-General Programme (2023A1515012319), and HKU-Seed Fund for Basic Research (202107185016, 2202100782). Thank Dr. Philip Wing-Lok Ho (Division of Neurology, Department of Medicine, The University of Hong Kong) for generously providing us with the mouse Neuro2a cell line.

## Author contributions

B.Y.L and L.L.G designed the study, carried out the research, analyzed, and interpreted the results and wrote the manuscript. X.Y.Y, and Z.Y.Z carried out the research and data analysis. S.L.C and L.W helped with the analysis of metabolites. L.G.J helped with the molecular, cellular, and biochemical experiments. R.C.L.H and W.J advised the study and edited the manuscript. All authors proofread the final manuscript and approved its submission.

## Competing interests

The authors declare no competing interests.

## Notes

### Competing Interest Statement

The authors have declared no competing interest.

